# *Populus* ERF85 balances xylem cell expansion and secondary cell wall formation in hybrid aspen

**DOI:** 10.1101/2021.06.08.447517

**Authors:** Carolin Seyfferth, Bernard A. Wessels, Jorma Vahala, Jaakko Kangasjärvi, Nicolas Delhomme, Torgeir R. Hvidsten, Hannele Tuominen, Judith Lundberg-Felten

## Abstract

Secondary growth relies on precise and specialized transcriptional networks that determine cell division, differentiation, and maturation of xylem cells. We identify a novel role for the ethylene induced *Populus ETHYLENE RESPONSE FACTOR ERF85* (Potri.015G023200) in balancing xylem cell expansion and secondary cell wall (SCW) formation in hybrid aspen (*Populus tremula x tremuloides*). Expression of *ERF85* is high in phloem and cambium cells and during expansion of xylem cells, while it is low in maturing xylem tissue. Extending *ERF85* expression into SCW forming zones of woody tissues through ectopic expression reduced wood density and SCW thickness of xylem fibers but increased fiber diameter. Xylem transcriptomes from the transgenic trees revealed transcriptional induction of genes involved in cell expansion, translation and growth. Expression of genes associated with plant vascular development and biosynthesis of SCW chemical components such as xylan and lignin, was downregulated in the transgenic trees. Our results suggest that ERF85 activates genes related with xylem cell expansion, while preventing transcriptional activation of genes related to SCW formation. The importance of precise spatial expression of *ERF85* during wood development together with the observed phenotypes in response to ectopic *ERF85* expression suggests that ERF85 functions as a switch between different phases of xylem differentiation during wood formation.

## 1. Introduction

Secondary xylem (wood) cells undergo cell expansion, extensive secondary cell wall (SCW) thickening and ultimately programmed cell death. SCW formation involves deposition of cellulose, xylan, and lignin (reviewed in [1, 2]). Understanding the molecular regulation of SCW formation in xylem vessels and fibers has become of special interest in tree research for breeding approaches and for generating a toolbox to tailor wood biomass production. High-resolution transcriptomic analysis of cambial tissues and developing xylem of aspen (*Populus tremula*) stems revealed transcriptional networks and putative molecular regulators of the different phases of xylem differentiation [3] that can be targets for reverse genetics. Functional analyses have furthermore revealed a set of WRKY, NAC and MYB transcription factors (TFs) as regulators of SCW biosynthesis [4–12].

The plant hormone ethylene has recently been shown to impact SCW formation in woody tissues. Gravitropic and exogenous stimulation of ethylene signalling leads to enhanced cambial activity, increased fiber-to-vessel ratio and to the formation of cellulose-rich gelatinous layers (G-layers) in xylem fibers, while these responses were abolished in transgenic ethylene insensitive hybrid aspen trees [13, 14]. Ethylene regulated TFs from the *PtEIN3 (ETHYLENE INSENSITIVE*) and the *PtERF (ETHYLENE RESPONSE FACTOR*) family are potential transcriptional regulators of SCW development in aspen [15–18], computational network analysis predicted *PtEIN3D* and eleven *PtERFs* as hub TFs in transcriptional regulation during stem development in *P. tremula* [17]. These TFs are co-expressed with genes related to lignan and xylan biosynthesis [17]. Furthermore, results from transgenic hybrid aspen (*P. tremula x tremuloides*) trees that overexpress PtERF139, or a dominant negative version of this TF, suggest a regulatory function of PtERF139 for SCW deposition and lignin biosynthesis [18]. In addition, ectopic expression of four other PtERFs affected stem diameter and wood biochemistry in hybrid aspen trees, indicating a functional link between PtERFs and wood formation [15]. One of the PtERFs identified in the aforementioned study was *ERF85*. Expression of *ERF85* is enhanced in hybrid aspen stems after ethylene treatment [15], but not in ethylene-insensitive hybrid aspen stems [14], suggesting that its function is controlled by ethylene. The closest homolog of *ERF85* in *Arabidopsis thaliana*, *AtCRF4* (*AT4G27950*), belongs to an evolutionarily distinct subgroup of *ERFs* called cytokinin response factors (*CRFs;* [19]). All *CRFs*, except *AtCRF4*, can be induced by exogenous cytokinin application [20]. The nine members of this subgroup can form homo- and heterodimers [21], suggesting that they might function together in gene regulation. Indeed, knock-out or overexpression of AtCRF4 alone in Arabidopsis resulted in a wild-type like phenotype under normal growth conditions, further supporting the idea of partial functional redundancy among the CRFs [22, 23]. On the contrary, ectopic expression of ERF85 alone was sufficient to modify tree growth and xylem cell wall chemistry [15].

In this study, we show how ectopic expression of ERF85 alters xylem formation by decrypting the biological processes and gene targets downstream of ERF85 in xylem tissues. For this, a detailed phenotypic characterization of the transgenic ERF85 trees (ERF85OE) was combined with a large-scale xylem transcriptome analysis using RNA-sequencing.

## 2. Materials and Methods

### 2.1 Plant growth conditions

Hybrid aspen (*Populus tremula L. x P. tremuloides* Michx, clone T89) was used in all experiments, propagated *in vitro* and transferred to soil. Greenhouse growth conditions were as follows; 18:6h day:night cycle, 20:15°C, day:night temperature, and relative humidity ranging between 50 and 60%. All trees were grown in a commercially available sand/soil/fertilizer mix (Krukjord, Hasselfors Garden, Örebro, Sweden) and fertilized once per week with approx. 150mL 1 % Rika-S (N/P/K 7:1:5; Weibulls Horto, Hammenhög, Sweden) starting the third week after transplanting and ending one week before harvest. Trees were rotated weekly to minimize positional effects. Growth conditions for trees used for wood density and anatomy analysis were slightly different: trees were grown under greenhouse conditions in a mixture of peat, sand and Vermiculite (6:2:1, v/v/v) and fertilized with 2.5g/L (w/v) of Osmocote Exact Hi-Start (N/P/K 15:4:8; Scotts). Temperature was 20/18°C (light/dark) and photoperiod 18h. Natural daylight intensity (on average 400μmol/ms) was controlled with curtains and supplemented with high-pressure sodium lamps when needed.

### 2.2 Cloning and generation of transgenic trees

To generate the *pERF85*::*GUS* construct a 1951-bp promoter region upstream of *ERF85* (Potri.015G023200) was amplified. Primers used can be found in Supplement Table S1. The PCR product was cloned into the pDONR221 donor vector and recombined to pKGWFS7, driving eGFP and GUS expression [24], thereafter cloned into Agrobacterium strain GV3101 pMP90 and transformed into hybrid aspen [25]. At least 17 independent transgenic lines carrying the *pERF85*:GUS construct were generated and selected following detection of GUS expression in stems, leaves and roots of *in vitro* grown trees.

### 2.3 Histochemical staining

GUS staining in wood material from greenhouse grown transgenic hybrid aspen trees expressing the *pERF85*::*GUS* construct was performed in three transgenic lines. Trees were grown in the greenhouse to a height of approximately 2m. Stem segments from 10cm above soil level were used for histochemical staining. For the promoter activity analysis in response to ACC, the transgenic lines have been grown in an *in vitro* culture system for four weeks. A ten-hour treatment of either 100μM ACC or mock (water) was applied (according to [14]). Three stem pieces from the fifth internode down were harvested and the sections shown are representing a region in the upper end of the lower third of the stem. Plant material was fixed in 90% acetone for 20min at room temperature. Acetone was dried off and samples were immersed in the GUS-staining solution containing 1mM X-GlcA (5-bromo-4-chloro-3-idolyl glucuronide), 50mM K-phosphate buffer (pH 7.0), 0.1% Triton, 1mM potassium ferricyanide (K_3_Fe(CN)_6_) and 1mM potassium ferrocyanide (K_4_Fe(CN)_6_). Samples were vacuum infiltrated for 3 cycles of 2min, followed by an incubation at 37°C in the dark and terminated after 6h (for greenhouse-grown trees) or 16h (for ACC-treated *in vitro*-grown trees). Samples were rinsed with phosphate buffer, cleared using an ethanol washing series and incubation in 100% ethanol over-night. Afterwards an ethanol washing series was used to rehydrate the samples. GUS expression was analyzed in 60-70μm thick stem cross-sections (cut using a vibratome) that were mounted in 50% glycerol and imaged with a Zeiss Axioplan2 microscope, Axiocam HRc camera and Axiovision V4.8.2 software (Carl Zeiss Light Microscopy, Göttingen, Germany). To check the start of SCW formation, stem cross sections (35μm) were made using the vibratome and stained with Safranin:Alcian blue (1:2 ratio) for one minute. Sections were rinsed twice in water and mounted in 50% glycerol.

### 2.4 Wood density measurements

A 20cm piece of stem (n=5) from (approximately) 2m tall tree was cut 10cm above the soil and oven-dried for one week at 65°C. Wood density was determined as the dry weight of wood sample divided by dry volume. The diameter of each stem sample was measured (with a calliper) from different positions (top, middle and bottom of the stem piece) three times. Each stem piece was still measured independently three times (as technical replicates) to ensure the correct volume.

### 2.5 Transmission electron microscopy

Approximately 0.5mm thick transverse hand-cut sections from greenhouse-grown stem pieces harvested 8-10cm above soil level were fixed in 2.5% glutaraldehyde in sodium cacodylate buffer (0.1M, pH7.2) for more than 48h, treated for 2h with 1% OsO4 and then processed, embedded, sectioned and imaged as described in [26].

### 2.6 Quantification of xylem cells, fiber diameter and cell wall thickness

Ultrathin stem sections (1μm thick) were stained with toluidine blue and pictures were taken with a 20-fold and 100-fold magnifying objective of a Zeiss Axioplan 2 microscope at approximately 1mm inwards the cambium. Each picture covers an area of approximately 3.02mm^2^ or 149,295μm^2^, respectively. Numbers of vessels and ray cells were counted in ImageJ using a picture taken with 20-fold magnification. The same pictures were used to quantify vessel diameter, defined as the area surrounded by the vessel cell wall. Pictures taken with 100-fold magnifying objective were used to quantify fiber number, diameter and lumen to lumen distance (from here on defined as SCW thickness). Sections from three trees per genotype were analyzed and fiber parameters were quantified on five different positions per section (within an area located approximately 1mm inwards the cambium). The dark-blue stained middle lamella was used to define the boundary of each fiber cell, and the area included by it was defined as fiber cell diameter. Cell wall thickness, defined as distance between the lumen of two neighbouring cells, was measured at 50 randomly chosen spots per picture. To calculate statistical differences between fiber numbers, diameter and cell wall thickness in wild-type (WT) and the independent transgenic lines, measurements from all five pictures taken from each tree were summarized to calculate the mean value.

### 2.7 RNA extraction

RNA was extracted from 15cm long stem pieces taken approximately 15-30cm above soil. Prior to RNA extraction, bark and pith of each stem piece were removed and the developing and mature xylem (cells with high expression of *ERF85* in ERF85OE) was scraped all around the stem and ground in liquid nitrogen with mortar and pestle. RNA was extracted from three WT samples (pooled from three trees each) and three samples (each a pool of three trees) for each line of the ERF85OE. RNA was used for RNA-Sequencing (RNA-Seq) and real-time quantitative PCR (qPCR). RNA extraction was performed using the BioRad Aurum RNA extraction kit following the manufacturer’s instructions, except for the on-column DNase treatment which was replaced by a DNase treatment using DNAfreeTM (Ambion) after RNA elution from the column. RNA was cleaned using the Qiagen MinElute kit. RNA was quantified using a Nanodrop ND-1000 (Nano-Drop Technologies, Wilmington, DE, USA) and quality was assessed with an Agilent 2100 Bioanalyzer (Agilent, Waldbronn, Germany) with Agilent RNA 6000 Nano Chips according to the manufacturer’s instructions.

### 2.8 RNA-Seq sample preparation and statistical analysis

Sequencing library generation and paired-end (2x 100bp) sequencing using Illumina HiSeq 2000 were carried out at the SciLifeLab (Science for Life Laboratory, Stockholm, Sweden). Raw files can be downloaded from the European Nucleotide Archive (ENA) under PRJEB35743. Data quality summary and scripts used in this study are available in the GitHub repository https://github.com/carSeyff/ERF85OE (DOI: 10.5281/zenodo.4770645), where details on non-default parameters for data pre-processing can be found. Briefly, data pre-processing was performed as described in Delhomme *et al*. (2014) and consisted of removal of ribosomal RNAs and sequencing adapters (using SortMeRNA (v2.1b using the rRNA libraries shipped with the tool) and Trimmomatic (v0.32) [27, 28]), as well as quality-based trimming by the latter. The *P. trichocarpa* (Pt) genome (version 3.0) was retrieved from http://popgenie.org and used for the alignment of the quality-filtered and trimmed read pairs using STAR (v2.4.2a). Read counting per gene and library was performed using HTSeq that discard ambiguous and multi-mapping reads [29]. Statistical data analysis was done in R (version 3.4.2) using DESeq2 (version 1.22.2 [30]). This analysis included library size adjustment to obtain the size factor value for each genotype (WT against ERF85OE lines) and variance stabilizing transformation (VST). The resulting normalized read counts were used for principal component analysis. The differential gene expression analysis between WT and the three lines of the ERF85OE was conducted on the raw counts. Differentially expressed genes (DEGs) were selected for ERF85OE compared to wild-type and defined by a |log2FC|>1 and a pAdj<0.01 (given in Supplementary Table 2). Heatmaps were generated in R with the pheatmap package (version 1.0.12) and gplots. All given Populus gene annotations are reported according to the gene name given to its closest *A. thaliana* homolog.

### 2.9 Validation of RNA-Seq results by qPCR

RNA used to validate gene expression changes in ERF85OE lines was extracted from three replicates per genotype and three pools of WT trees. cDNA was synthesized from 1μg RNA using iScript cDNA Synthesis Kit (BioRad). Each qPCR reaction (in total 15μl) contained 2μl of diluted cDNA (1:4), 7.5μl 2xSYBR Green Mastermix (Roche) and 2.75μl of 10nM forward and reverse primers (listed in Supplement Table S1) and run in the Bio-RAD CFX96 Real Time System. Target gene expression was compared with expression of three reference genes (PtACT1 and PtUBQ-L), which showed stable expression pattern in poplar stems (Wang et al., 2016). PtUBQ-L was chosen as reference gene for calculating gene expression changes (presented as log2 of ΔcT) shown in Figure 4D.

### 2.10 Bioinformatic and statistical analysis

Gene ontology and PFAM enrichment were performed using *P. trichocarpa* homologs in AspWood based on Blast2Go [3]. Statistical analysis of phenotypic data was performed in R using one-way ANOVA and post-hoc HSD Tukey test. Each experiment was replicated using at least three trees per transgenic line. Multiple comparison tests were done using the R package multcompView (version 0.1-7), using a cut-off of p-value<0.05. Promoter motif analysis was done as described in [14]. In brief, promoter regions with different lengths (500bp, 1kbp, 1.5kbp and 2kbp) were obtained from the *P. trichocarpa* genome using the webtool “sequence search” from Popgenie (http://popgenie.org/sequence_search). We were able to obtain promoter regions of 401 (down-regulated) and 1837 (up-regulated) genes and checked the occurrence of the GCC-Box (“AGCCGCC”), the ERF-Box (“GCCGCC”) or the DRE-box (“RCCGAD”) on both strands. The hypergeometric distribution was used to determine statistical enrichment of either of the motifs among down- or up-regulated genes.

## 3. Results

### 3.1 ERF85 is expressed in the cambium and expanding xylem

The *P. trichocarpa* (Pt) genome encodes 170 *ERF*s, subdivided into ten different clusters [15]. The *CYTOKININ RESPONSE FACTORS* (*CRFs*) form a cluster in the *ERF* gene family (cluster VI; PtERF81-PtERF88) that comprises homologs to regulators of root, shoot, leaf and flower development identified in Arabidopsis [19, 23, 31]. Phylogenetic analyses identified eight *CRFs* in *P. trichocarpa* ([15]; Figure 1(a)). A high-resolution expression atlas derived from longitudinal sections through aspen stems [3], from here on referred to as the “AspWood” database, indicated high transcript abundances for all *PtCRF4* family members in samples representing cambial cell proliferation and xylem cell expansion (Figure 1(a,b); Supplement Figure S1(a)). Transcript levels of *ERF85* are high in the phloem and vascular cambium, peak in the xylem expansion zone and drop towards xylem maturation and secondary cell wall (SCW) formation before slightly increasing in the cell death zone. As ethylene is known to interfere with cambial activity and cell expansion [14], and high *PtCRF* expression was observed in the corresponding developmental zones, we explored the responsiveness of *PtCRFs* towards the ethylene precursor ACC (1-aminocyclopropane-1-carboxylic acid) in stems of wild-type trees and two ethylene-insensitive trees in published datasets (Figure 1(c); [14]). Treatment with the ethylene precursor ACC for 10h triggered induction of *PtERF81*, *PtERF82* and *ERF85* expression, and down-regulated expression of *PtERF86* (Figure 1(c)) in stems of wild-type but not ethylene insensitive trees, suggesting that these ERFs operate downstream of the ethylene signalling pathway. Phylogenetic analysis indicated 80% sequence identity between *ERF85* and *PtERF86* [15]. The high sequence similarity is typical for a gene duplication event and is often associated with redundant protein function. However, their opposite responses to ACC suggests that, despite high sequence similarity, regulation and potentially also function of *ERF85* and *PtERF86* might differ during ethylene signalling. This prompted us to focus our further analysis only on the role of *ERF85* in ethylene-regulated wood development. We examined the expression of *ERF85* in stem tissues of transgenic hybrid aspen (*P. tremula x tremuloides*) trees carrying a *GUS* reporter gene under the control of the 1951-bp *ERF85* promoter (Figure 1(d, e)). GUS staining confirmed the expression pattern obtained from AspWood, showing expression of *ERF85* mainly in phloem and expanding xylem cells, but also in contact ray cells next to vessel elements.

**Figure 1.**
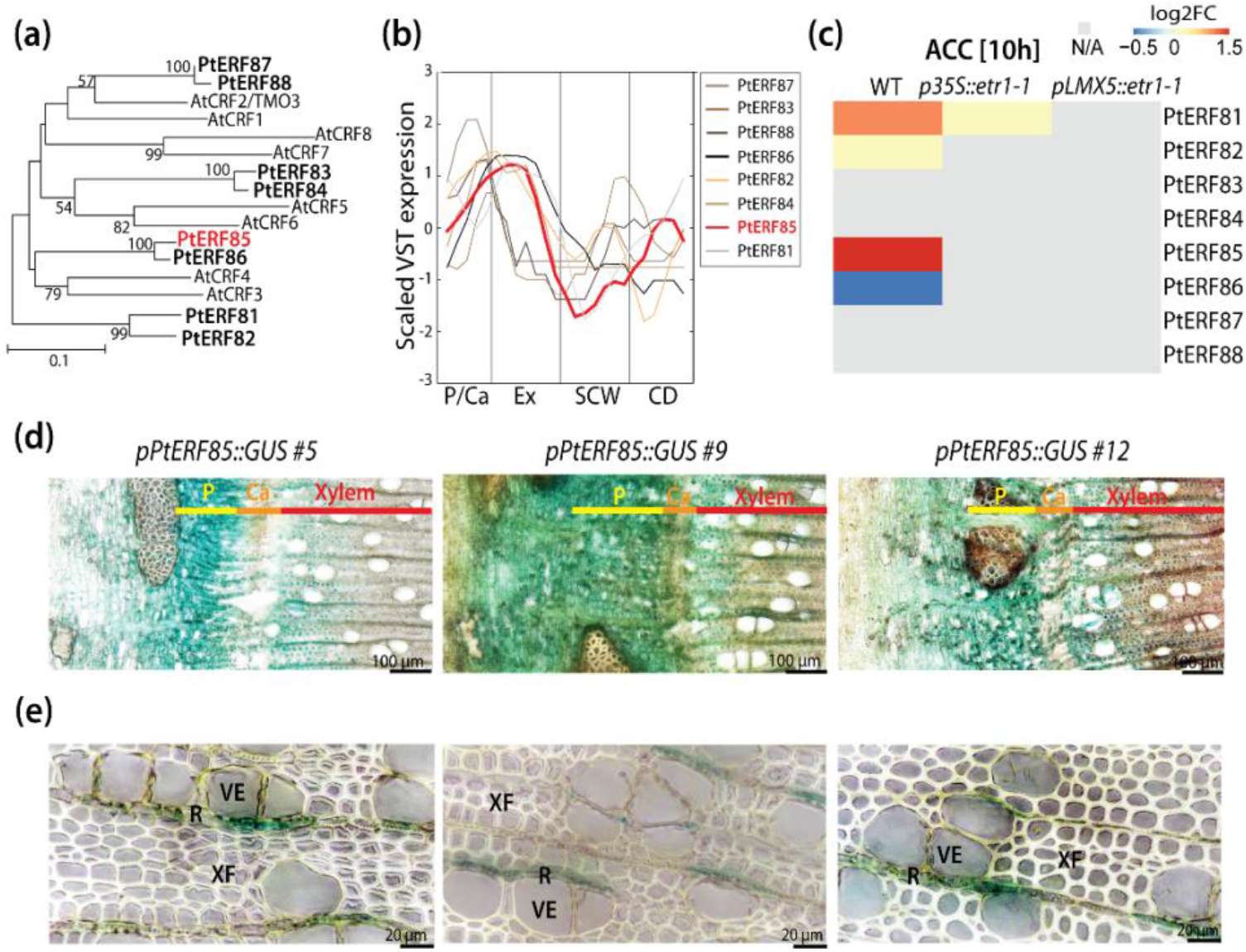
Expression of *ERF85* is observed during cambial growth and expansion of xylem cells. **(a)** Phylogenetic tree of *PtCRFs* (cluster VI; *PtERF81-PtERF88*) and its homologs in *A. thaliana* (*At*) (modified from [15]). **(b)** Expression profile of *PtCRFs* in aspen stems. Data (shown for one representative tree (T1)) was obtained from the transcriptome atlas AspWood [3]. To emphasize expression differences, scaled and smoothened VST expression values (implemented function in http://popgenie.org) are shown. **(c)** Heatmap showing the expression pattern of *PtCRFs* (log2 fold changes) in xylem tissue of wild-type (WT, clone T89) and two ethylene-insensitive trees (*p35S*::*etr1-1*, *pLMX5*::*etr1-1*) in response to 10h treatment with 10μM ACC (data extracted from [14]). **(d-e)** Cross sections of greenhouse grown trees showing GUS activity in three transgenic hybrid aspen trees expressing pERF85::GUS reporter gene. P=phloem; Ca=cambium; Ex=expanding xylem; SCW=secondary cell wall formation; XF=xylem fiber; VE=vessel element; R=ray. Gene names refer to *ERF81* (*Potri.019G131300*), *ERF82* (*Potri.013G158500*), *ERF83* (*Potri.002G167400*), *ERF84* (*Potri.014G094500*), *ERF85* (*Potri.015G023200*), *ERF86* (*Potri.012G032900*), *ERF87* (*Potri.001G094800*), *ERF88* (*Potri.003G136300*).

### 3.2 Ectopic ERF85 induction does not change ethylene response in hybrid aspen wood

The observed induction of *ERF85* expression by ethylene/ACC [15] prompted us to further investigate the role of *ERF85* in the ethylene signalling pathway. GUS reporter assays suggest that the stem expression pattern of *ERF85* was not altered by a 10 hours 100μM ACC treatment and remained concentrated predominantly to the cambium and the xylem (Figure 2(a)). It should be noted that, compared to *pERF85:GUS* expression in greenhouse-grown trees (Fig. 1(d, e)), GUS staining was faint in stems of *in vitro* grown trees (Figure 2(a)), making quantification of any response impossible. To study the function of *ERF85* in ethylene-regulated secondary growth, we examined the effect of ACC on transgenic stems of hybrid aspen trees that ectopically express *ERF85* under the xylem-specific *pLMX5* promoter (“ERF85OE”, described in [13]). According to AspWood (Supplement Figure S1(b)) and previous promoter GUS activity studies [13], expression of *LMX5* is highest during SCW formation. Wild-type trees showed only very low *ERF85* expression during SCW formation (Figure 1(e); Supplement Figure 1(b)). Using *pLMX5* therefore extends and increases *ERF85* expression beyond the cambium and xylem cell expansion zone into the SCW formation zone (Supplement Figure S1), thus spanning all zones that are affected by ethylene application [13, 14]. Upon ACC treatment, wild-type trees showed enhanced cambial activity, reduced vessel frequency and the induction of gelatinous-layer (G-layer) in xylem fibers [13, 14]. These phenotypes were not observed in ethylene-insensitive trees, which express a dominant negative version of the ethylene receptor ETR1 driven by the *pLMX5* promoter [13], indicating that ACC-induced xylem growth and G-layer formation requires functional ethylene signalling. In ERF85OE trees, ACC induced xylem growth and G-layer formation and favored fiber over vessel formation similarly to wild-type trees (Figure 2(b)). We detected patchy G-fiber occurence even in water-treated control trees (indicated by an arrow), which can randomly occur in fast growing *Populus* trees. We did not confirm a consistent G-fiber induction in ERF85OE trees in additional experiments where we investigated wood anatomy of greenhouse grown plants (Figure 3(b)). ACC treatment also reduced stem height growth of ERF85OE lines (Figure 2(c)), another phenotypic response seen in response to ACC application to *Populus* trees [13]. Together with previous results [15], this indicates that, even though *ERF85* can be transcriptionally induced by ethylene in an ethylene-signalling dependent way, it does not mimick constituitive ethylene signalling when ectopically expressed under *pLMX5* in woody tissues and still allows for normal ethylene responses.

**Figure 2.**
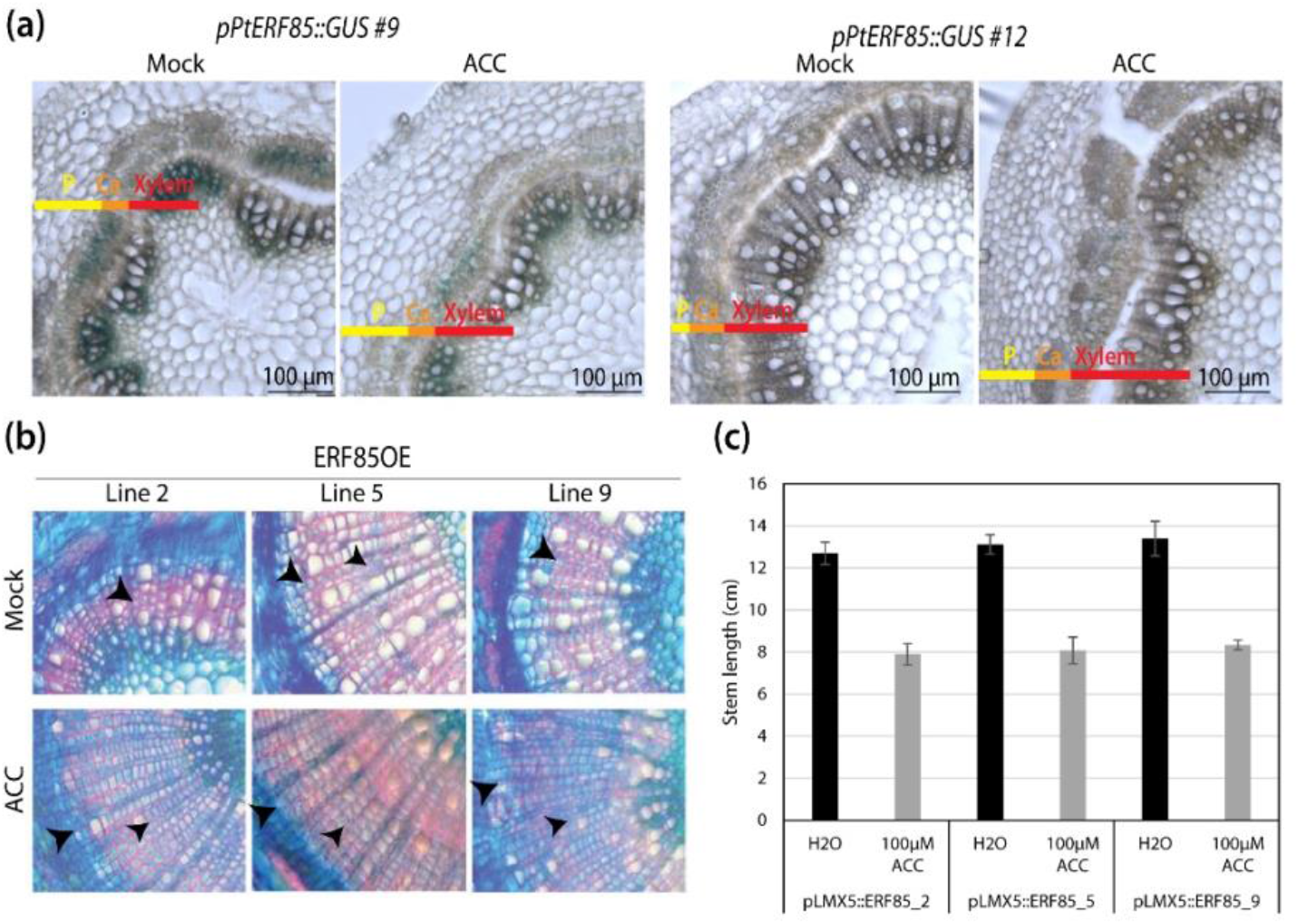
ERF85OE lines show enhanced ethylene-mediated stem phenotypes. **(a)** GUS staining in two transgenic lines that carry a *pERF85*::*GUS* reporter. Transgenic reporter lines were treated for 10h with either mock (water) or 100μM ACC. **(b)** Stem cross sections from WT, transgenic ethylene insensitive trees (pLMX5::etr1-1; [13]) and ERF85OE lines. Trees were treated with 10μM ACC for 14 days. Cross sections (50 μm thick) were stained with safranine and toluidine blue and magnified using a 20-fold objective. Arrows highlight the onset of a gelatinous (G)-layer in xylem fibers [14]. **(c)** Plant height of WT, ethylene-insensitive trees and ERF85OE lines in response to water (mock) or 100μm ACC treatment for 14 days. Bars represent the average height (+/-standard error) obtained from five biological replicates.

### 3.3 ERF85OE lines show increased xylem fiber diameter

In order to understand how the expansion of the expression zone of *ERF85* under the *pLMX5* promoter would affect xylem formation, we undertook a detailed analysis of xylem anatomy and subsequently assessed its downstream targets during wood development (section 3.4). In agreement with previous experiments, two out of three ERF85OE lines analyzed showed a reduced stem growth (Figure 3a; [15]). Observation of ultrathin transverse sections either through transmission electron microscopy or, after toluidine blue staining, through light microscopy, suggested changes in xylem fiber morphology (Figure 3(b)). Secondary xylem anatomy of the ERF85OE lines was analyzed in xylem areas with clearly visible SCWs, approximately one mm radially inwards of the cambium (Supplement Figure S2(a)). The average number of fibers (per 149 295 μm^2^ area) was reduced in all transgenic lines, up to 16% (line 5) compared to wild-type (Figure 3(c)). No significant differences (p-value<0.05) were observed in the total number of ray or vessel cells between wild-type and any of the ERF85OE lines (Figure 3(d-e), covering an area of 3.02mm^2^ shown in Supplement Figure S2(b)). The fact that the number of fibers in a defined cross-sectional area of the secondary xylem was reduced in the ERF85OE lines, but the numbers of the ray cells or vessels were not altered, suggests an increase in fiber diameter. Indeed, fiber diameter, defined in the transverse sections as the area encircled by the middle lamella, was increased by up to 27% (line 5) (Figure 3(f)). No consistent change was observed in vessel diameter (Supplement Figure S2(c)). We next analyzed the effect of ectopic *ERF85* expression on cell wall and mechanical properties. Using transmission electron microscopy, we observed that fibers had thinner cell walls compared to wild-type (Figure 3(g)). Quantification of fiber cell wall thickness (defined as distance between cell to cell lumen) on toluidine blue stained ultrathin stem cross sections showed a reduction (between 10-19%) in cell wall thickness for ERF85OE lines compared to wild-type (Figure 3(g)). Furthermore, ERF85OE trees showed a reduction in wood density, ranging from 8 to 12% compared to wild-type trees (Figure 3(h)). The decrease in xylem density in ERF85OE lines is therefore likely the result of thinner fiber walls and an increased fiber diameter. Taken together, ectopic expression of *ERF85* throughout all phases of wood development results in increased fiber expansion and reduced SCW deposition, and ultimately in reduced wood density.

**Figure 3.**
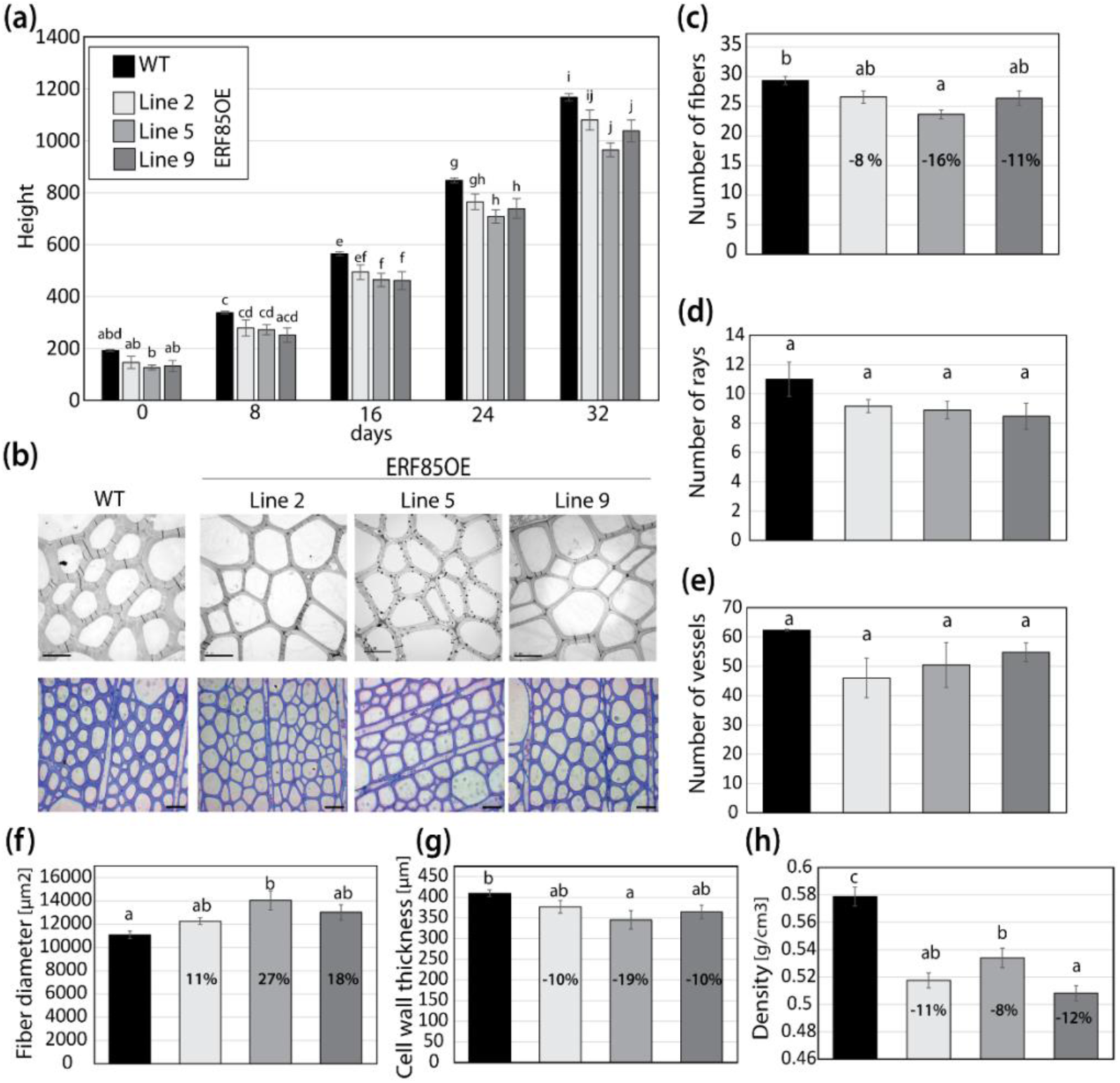
Ectopic expression of *ERF85* in woody tissue increases the size of the fibers and reduces cell wall thickening. **(a)** Stem growth rate of wild-type (WT) and three transgenic lines that ectopically express ERF85 (ERF85OE) under a xylem specific wood promoter (*pLMX5*; [15]) in controlled greenhouse environment. **(b)** Transmission electron micrographs (TEM; 100-fold maginification; scale bar is 10μm) and toluidine blue staining (20-fold maginification; scale bar is 40 μm). TEM pictures cover a total area of 149295μm2. **(c)** The number of fiber cells counted in toluidine blue cross-sections shown in (b). **(d-e)** Numbers of ray and vessel cells (in an area of 3.02mm2) in WT and ERF85OE lines. Cross-sections used for quantification are shown in Supplement Figure S2(b). **f)** Fiber diameter from toluidine blue-stained cross-sections shown in (b). Cell outlines were defined by the middle lamella and the enclosed area [μm2] was measured using ImageJ. **(g)** Cell wall thickness (determined as lumen to lumen distance of two neighbouring cells [μm]) based on toluidine blue-stained stem sections (shown in (b)), using a 100-fold magnification. **(h)** Wood density. For all panels, bars represent mean ± SE calculated from three biological replicates per line using a linear effect model. Significant differences between genotypes are indicated by unique occurrence of letters above the bars and were assigned based multiple comparison test and a p-value<0.05.

### 3.4 *Ectopic expression of* ERF85 *activates expression of genes linked to cell expansion and antagonizes induction of genes related to SCW formation*

To understand the molecular mechanisms underlying *ERF85* function in xylem differentiation, transcriptome data was generated by RNA-Seq from developing xylem of wild-type trees and the ERF85OE lines (Figure 4). Principal component analysis (PCA) on normalized gene count data revealed separation of the wild-type samples from the ERF85OE samples in the first component (explaining 61% of the overall variation), suggesting that genotypic differences explain most of the variation among the transcriptomes (Figure 4(a)). Differential analysis of genes showing at least a two-fold change between ERF85OE and wild-type (at a False Discovery Rate (FDR) cutoff of 1%; Supplement Table S2) identified 1893 genes with higher expression in ERF85OE compared to wild-type, while expression of 410 genes was down-regulated (Figure 4(b)). Transcripts down-regulated by ectopic expression of *ERF85* expression were enriched in GO terms related to metabolism and signalling (Supplement Figure S3(a); Supplement Table S3), including homologs with known function in xylem cell wall composition (Figure 4(c); e.g. *IRX10* (*IRREGULAR XYLEM10*), *4CL2* (*4-COUMARATE*:*CoA LIGASE2*)) and xylem development (e.g. *XCP2* (*XYLEM CYSTEINE PEPTIDASE2*), *CLV20* (*CLAVATA20*)). Expression of the *PIN3* homolog was also supressed in ERF85OE (Figure 4 (c,d)), which function in polar auxin transport has been linked to increased xylem formation in Arabidopsis [32]. DEGs with upregulated expression in ERF85OE were enriched (p-value<0.01) in Gene Ontology (GO) terms related to protein translation (n=335; Supplement Figure S3(b); Supplement Table S4) including genes encoding ribosome subunits, elongation factors, cyclophilins and chaperons. In addition, we also identified homologs with reported function in xylem development and cell elongation (Figure 4(c)), including the endo-1,4-ß-glucanase CEL that affects xylem development and cell wall thickness in Arabidopsis [33] and cell growth and cellulose content in suspension-cultured poplar [34]. The observed change in expression was validated for both, up- and down-regulated DEGs, using qPCR (Figure 4(d)). We next assessed the stem expression profile of the up-regulated DEGs in AspWood (Figure 4(e)). About 67% (1304 out of 1893 genes) of the DEGs are present in AspWood clusters that show, like ERF85, highest expression in the cambium and during xylem cell expansion, but low expression during SCW formation (clusters “f”, “e”, “h” in [3]). Approximately 33% of all down-regulated DEGs (137 out of 410 genes) belong to AspWood clusters dominated by genes with opposite expression profiles to *ERF85*, meaning highest expression during SCW formation and low expression in cambium and expansion zone (cluster “g” in [3]). Thus, *ERF85* might act positively on the expression of gene networks underlying secondary xylem cell expansion, while it has a negative effect on the onset of gene expression in networks that underly SCW formation.

**Figure 4.**
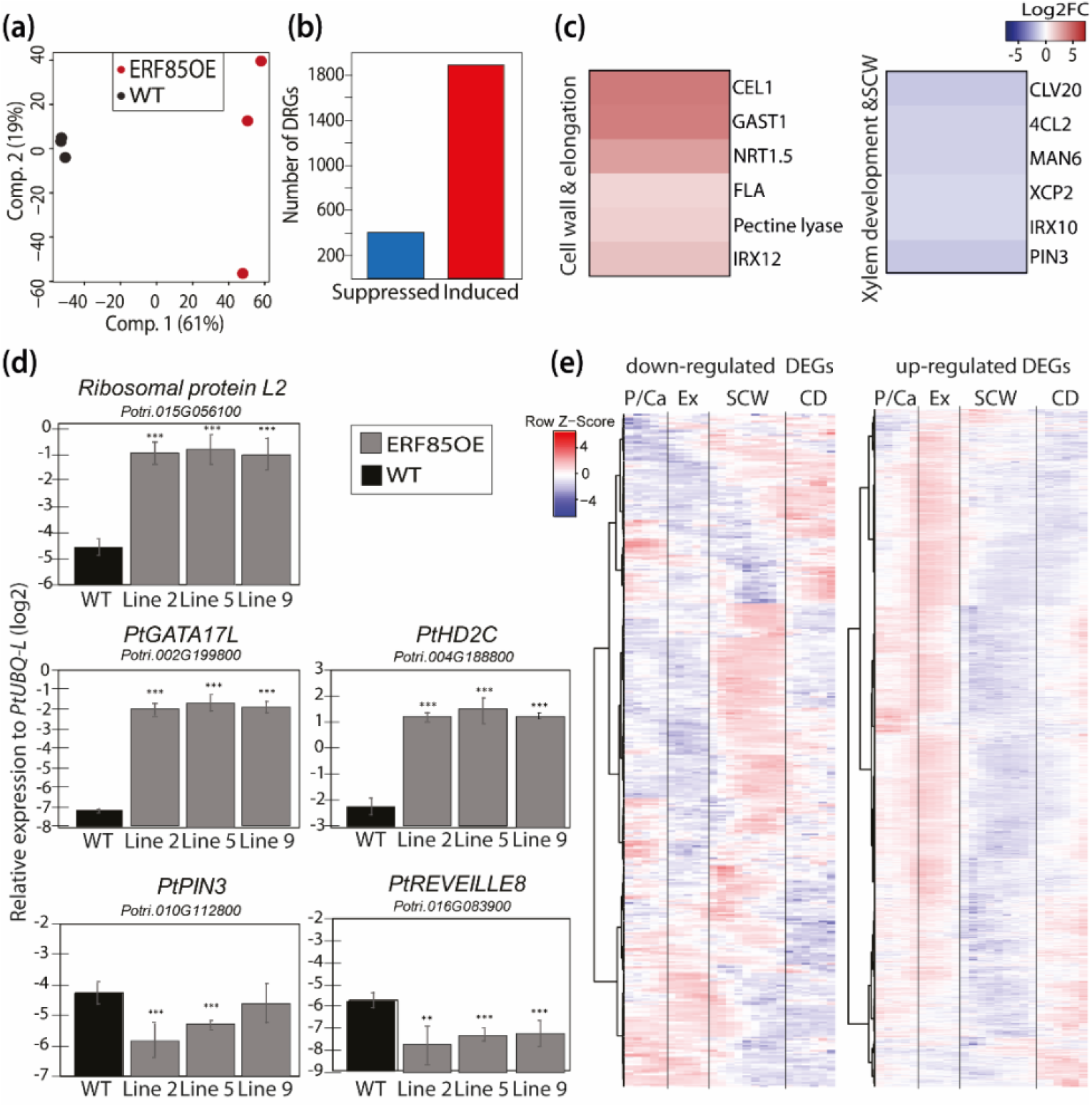
*ERF85* stimulates expression of genes linked to cell expansion and supresses genes linked to secondary cell wall formation during xylem differentiation. **(a)** Principal component analysis of xylem transcriptome data obtained from three wild-type samples (WT, each sample represents a pool of three plants) and three ERF85OE lines (2,5,9). **(b)** Distribution of up- and down-regulated differentially regulated genes (DEGs) in ERF85OE. DEGs were selected based on a |log2FC|>1 and pAdj< 0.01 compared to WT. **(c)** Representative genes found among the DEGs in ERF85OE lines with known function of their homologs in Arabidopsis related to cell elongation, cell wall composition and xylem development (*Potri.015G127900* (*CEL1*); *Potri.002G022600* (*GAST1*); *Potri.001G171900* (*Pectin-lyase*); *Potri.008G064000* (*IRX12*); *Potri.014G156600* (*CLV20*); *Potri.010G112800* (*PIN3*); *Potri.005G256000* (*XCP2*)) and secondary cell wall (SCW) composition (*Potri.010G192300* (*FLA*); *Potri.001G036900* (*4CL2*); *Potri.016G138600* (*MAN6*); *Potri.005G256000* (*IRX10*)), *POPTR_0003s08710* (*NRT1.5*). **(d)** Validation of differential expression of up- and down-regulated DEGs using quantitative real-time PCR (qPCR). Bars represent mean + SD expression level of target genes relative to the housekeeping gene *PtUBQ-L* (log2). Asterisks indicate statistically significant differences between wild-type (WT) and ERF85OE lines according to Student T-Test (with p-value<0.05 (*), p-value<0.01 (**), and p-value<0.001 (***)). **(e)** Heatmap showing the expression profile of up- and down-regulated DEGs during stem development in hybrid aspen. Data was obtained from the AspWood database [3]. Genes are scaled so that red represent the highest expression for that gene, and blue the lowest, across all developmental zones. P/Ca=phloem/cambium, Ex=expanding xylem, SCW=secondary cell wall formation, CD=cell death.

We utilized the AspWood co-expression network to cluster DEGs according to their expression profile during xylem development (Figure 5). This captured 801 DEGs with high co-expression, which clustered into 13 gene modules (Figure 5(a); Supplement Table S5). According to their stem expression profiles and functional gene characterization we classified them into gene modules underlying xylem expansion (M1-M5), transitioning from primary to SCW (M6, M7), SCW formation (M9-M11) and fiber cell death or ray development (M8, M12, M13). Cell expansion-related modules M1-M4 comprised the highest number of DEGs with increased expression in ERF85OE and an ERF85-like stem expression profile (Figure 5(a, b)). These four expansion-related modules consisted of genes related to protein translation and folding, such as elongation factors (e.g. *EUKARYOTIC INITIATION FACTOR4A-2* (*PtEIF4A2*), RNA binding (e.g. *RNA-BINDING PROTEIN47C* (*PtRBP47C*)), rRNA processing and maturation (which is partially controlled by HISTONE DEACETYLASE2C (PtHD2C) [35]), ribosomal subunits (e.g. *RIBOSOMAL PROTEINS27* (*PtRS27A*)) and chaperones (e.g. *CHAPERONIN-60 BETA2* (*PtCPNB2*), Supplement Table S6). Module (M5) contained genes, which showed a peak in expression during xylem cell expansion but, in contrast to M1-4, remained high at the interface between cell expansion and the onset of SCW formation. This module included genes that encode pectin-degrading polygalacturonases (e.g. *PtPG2*, *PtPGX3*) and ATP-binding cassette transporter. Interestingly, *PtPGs* that were lowly expressed in the SCW zone in AspWood (e.g. *PtPG9*, *PtPG2*) were up-regulated in ERF85OE, whereas PtPGs that were highly expressed in the SCW zone in AspWood (e.g. *PtPG6*) were repressed in ERF85OE. Transcriptional reprogramming of pectin modulators could lead to loosening of pectin structures to facilitate cell expansion, as for example shown for the PtPG28/43 homolog in Arabidopsis *AtPGX3* [36], and a reduction of pectins in SCWs [37]. Also the two modules M6 and M7 (Figure 5(c)) contain genes which in AspWood show high expression during the transition phase from primary to SCW in AspWood, including genes involved in flowering (e.g. cyclin *DOF2* (*PtCDF2*)), response to light (e.g. *CRYPTOCHROME1* (*PtCRY1*)) and the circadian clock (e.g. *REVILLE8* (*PtRVE8*)), as well as the negative regulator of vessel formation *VND-INTERACTING2* (*PtVNI2*). The three SCW-associated modules (M9-M11, Figure 5D) contained genes associated with auxin signalling (e.g. *PtPIN3*, *PtIAA4*), biosynthesis of the SCW-specific hemicellulose xylan (e.g. *PtIRX10*) and lignin (e.g. *Pt4CL2*) and xylem maturation (e.g. *PtXCP2*). Expression of all genes in these three modules was down-regulated in ERF85OE, which agrees with the reduction of SCW formation in the transgenic trees (Figure 3(g)). Finally, three gene modules (M8, M12, M13, Figure 5(e)) showed highest expression in the cell death zone. While expression of genes in M8 and M12, such as *CHALCONE SYNTHASE (PtCHS*), which is involved in biosynthesis of flavonoid synthesis and can prevent auxin transport, was increased in ERF85OE, expression of genes in M13 (such as *AMMONIUM TRANSPORTER2 (PtAMT2*)), was down-regulated in ERF85OE. In agreement with increased fiber diameters in ERF85OE lines, we observed that ectopic expression of ERF85 throughout all stages of xylem development induces genes specific to xylem cell expansion. At the same time, genes with highest expression during SCW formation are down-regulated in ERF85OE. Thus, radially prolonged transcriptional activation of genes involved in xylem cell expansion, and the consequent indirect suppression of SCW-related gene expression could ultimately cause the observed reduction of fiber cell wall thickness.

**Figure 5.**
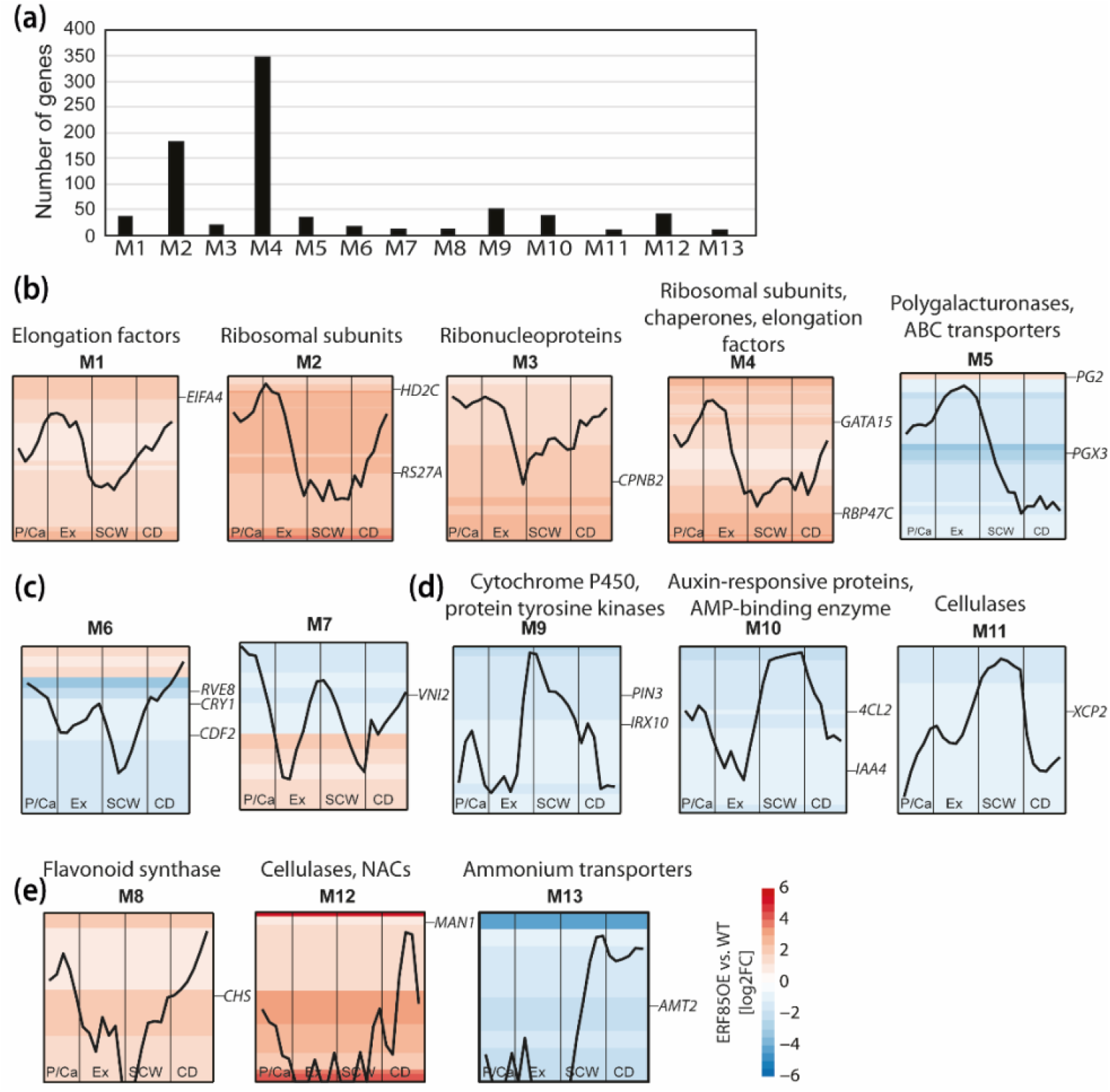
Ectopic expression of *ERF85* induces expression of genes related to translation, while suppressing expression of SCW-related genes. **(a)** Genes with similar expression profiles during stem development were categorized into 13 gene modules (M1-M13) based on the AspWood co-expression network [3]. The size of each gene module is given in the histogram. **(b)** Gene modules with highest expression during xylem cell expansion. A representative gene, defined as the gene with the highest degree, was chosen to illustrate the expression profile of genes in each module (Supplement Table S7). Enriched PFAM terms are listed above each module (Supplement Table S6). Background heatmaps indicate gene expression (log2FC) in ERF85OE compared to wild-type for the genes in the module (each row represents a gene). Significantly enriched PFAMs are listed above each module (p-value<0.05). **(c)** Gene modules with highest expression in the transition phase of primary to secondary cell wall formation. **(d)** Gene modules with highest expression during SCW formation. **(e)** Gene modules with highest expression in the zone of cell death. P/Ca=phloem/cambium; Ex=expanding xylem; SCW=secondary cell wall formation, CD=cell death; *EIFA4-2* (*Potri.001G197900*); *HD2C* (*Potri.004G188800*); *RS27A* (*Potri.003G161200*); *CPNB2* (*Potri.001G002500*); *GATA15* (*Potri.008G213900*); *RBP47C* (*Potri.002G128100*); *PG2* (*Potri.001G171900*); *PGX3* (*Potri.008G100500*); *RVE8* (*Potri.016G083900*); *CDF2* (*Potri.004G121800*); *CRY1* (*Potri.005G164700*); *VNI2* (*Potri.017G063300*); *CHS* (*Potri.003G176700*); *PIN3* (*Potri.010G112800*); *IRX10* (*Potri.003G162000*); *4CL2* (*Potri.001G036900*); *IAA4* (*Potri.010G078400*); *XCP2* (*Potri.005G256000*); *MAN1* (*Potri.001G266900*); *LBD4* (*Potri.007G066700*); *AMT2* (*Potri.019G000800*).

### 3.5 ERF85-*mediated transcriptional regulation mechanisms during xylem cell expansion*

We next compared *A. thaliana* homologs identified as DEGs in our dataset with direct targets of AtCRF4 in *A. thaliana* [38, 39]. Of the 2303 DEGs, 2227 had corresponding *A. thaliana* homologs. However, because of gene duplication events in *Populus*, 475 DEGs did not have unique *A. thaliana* homologs. In total we identified 1752 unique *A. thaliana* homologs among the initial 2303 DEGs. Dataset comparison to the targets of AtCRF4 revealed that 995 out of these 1752 *A. thaliana* homologs had been identified as direct AtCRF4 targets (Supplement Table S2). Their *Populus* homologs are therefore potential direct targets of *ERF85*. This includes both homologs of up- and down-regulated DEGs of ERF85OE (Figure 6(a)). These results point towards a direct regulation of DEGs by *ERF85*. In support of this, promoter regions (from 500 up to 2000bp upstream of the start codon) of up-regulated DEGs showed a significant enrichment of the ERF-binding motif GCC-Box (p-value<6.01*^e^-11^ and p-value<1.87*e^^-5^, for 0.5 and 2kbp, respectively). However, no statistically significant enrichment was identified for this motif in promoters of down-regulated DEGs (Supplement Table S8).To determine how *ERF85* could achieve transcriptional regulation during different developmental stages in xylem tissues, we analyzed co-expression relationships of the *ERF85*-regulated TFs during wood development (Figure 6(b), Supplement Table S9). *ERF85*-regulated TFs were grouped based on their expression profile in AspWood into TFs with low expression during SCW formation (e.g. *PtGATA15*, *PtHD2C*; Figure 6(b)) and TFs with high expression during SCW formation (e.g. *HEAT SHOCK TRANSCRIPTION FACTOR3* (*PtHSFB3*), *PtWRKY21*, *PtERF151*). The GCC-box (AGCCGCC) and the DRE-motif (GTCGGT/C) were present in promoters (two kb) of TFs from both groups (Supplement Table S9), suggesting a potential direct transcriptional regulation of certain targets by *ERF85*. Indeed, the closest homolog of ERF85 in *A. thaliana*, *AtCRF4*, controlled expression of 18 TF homologs (out of all 57 TFs identified in our study, p-value=0.17) by direct binding (highlighted in Figure 5(b), Supplement Table S2; [38]). Among them, we identified homologs of *PtHD2A* and *PtGATA17L*. We ranked each TF based on its network degree (number of neighbours) with the aim to select TFs with central regulatory function downstream of *ERF85*. Several PtGATAs (*PtGATA15*, *PtGATA16*, *PtGATA17L*) and two histone deacetylases (*PtHD2A*, *PtHD2C*), which we included because they regulate gene expression by promoter binding [35, 40, 41], showed a central position in the TF network downstream of *ERF85*. Their expression during wood formation follows the same pattern as ERF85 and although only PtHD2A and PtGATA17L were identified as direct targets in AtCRF4 [38], we identified the GCC-box (PtGATA17L) or DREB-motif (PtGATA15, PtHD2C) in their promoter region (Supplement Table S9). PtHD2A, PtHD2C and PtGATA17L have been identified in gene modules up-regulated during cell expansion (Figure 5(b)) and homologs of DEGs from ERF85OE were found as direct target genes of AtGATA17L and AtHD2C in *A. thaliana* (Figure 6(a), Supplement Table S9; [35, 39]). Furthermore, according to the AspWood co-expression network analysis (Figure 6(b); Supplement Table S5), expression of *PtGATAs* (like *PtGATA15* in gene module M4) and both, *PtHD2C* and *PtHD2A* (present in gene module M2), was linked to the expression of genes encoding ribosomal subunits, chaperones and elongation factors. These results point towards a central role of *PtGATA17L* and *PtHD2C* in transcriptional regulation of ribosome biogenesis and translation downstream of ERF85 during xylem cell expansion. Concurrently, *ERF85* could prevent the onset of SCW formation by suppressing central TFs in the underlying network, such as *PtHSFB3* or *PtERF151* (both containing a DRE-motif in their promoter and belonging to gene module M10; Supplement Table S9).

**Figure 6.**
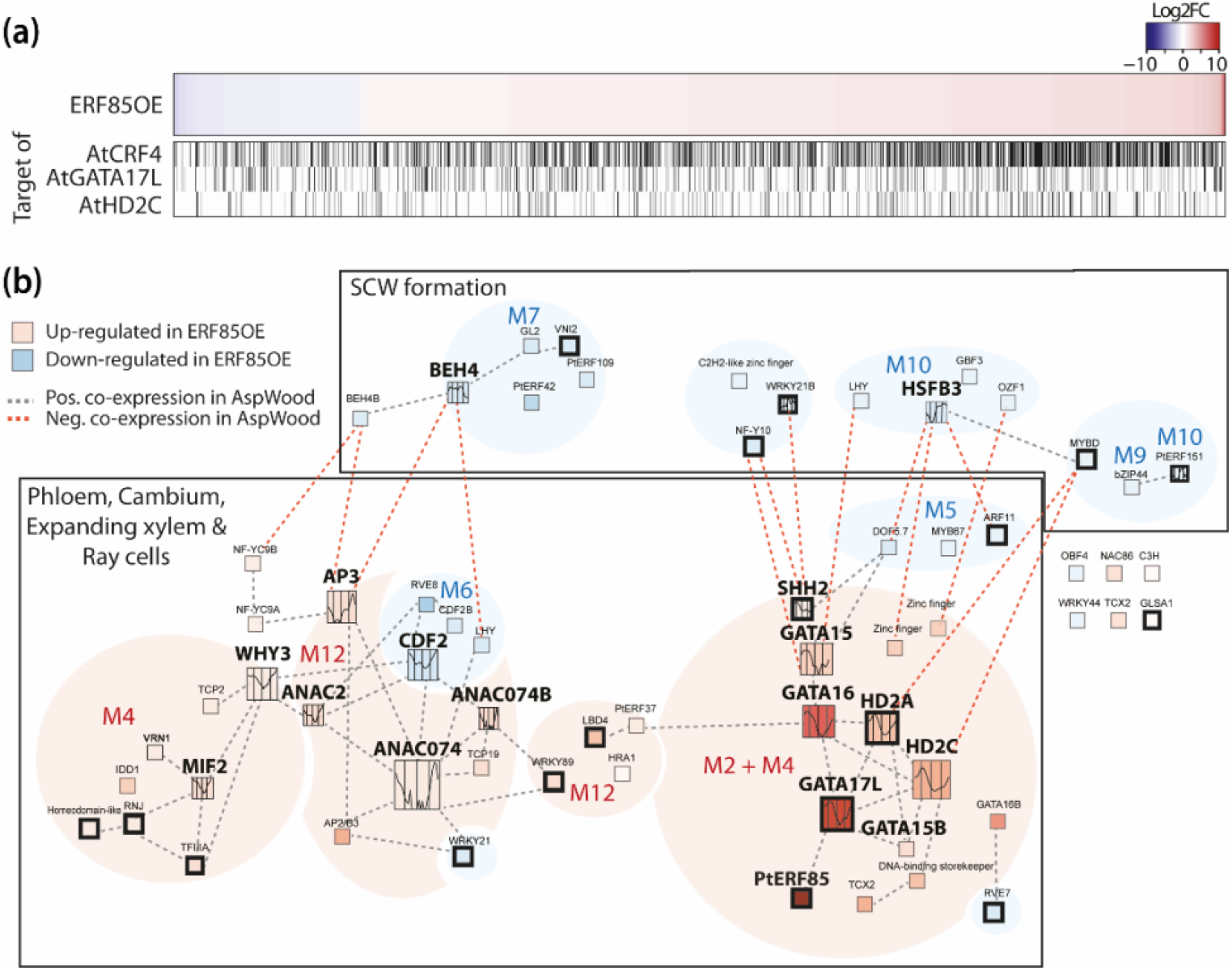
*ERF85*-controlled gene regulatory networks during wood development. **(a)** Heatmap showing gene expression of all DEGs in ERF85OE compared to wild-type. The graphs beneath the heatmap indicate homologs that were identified as direct targets of *AtCRF4*, *AtGATA17L* and *AtHD2C* (Supplement Table S2; [35, 38, 39]). **(b)** Co-expression network of differentially expressed TFs of ERF85OE during wood development (Supplement Table S9). Thickened box lines indicate TFs for which homologs were identified as direct targets of *AtCRF4* [38]. Background colour represents gene expression induction (red) or suppression (blue) in ERF85OE lines compared to wild-type. TFs with at least four neighbours were highlighted by increased box size and expression profiles inside the box show their expression pattern during wood formation (obtained from the AspWood data base [3]). Grey lines indicate positive co-expression between the TFs, red indicates negative co-expression. M=module (corresponding to Figure 5).

## 4. Discussion

### 4.1 ERF85 modulates the balance between xylem cell expansion and SCW formation through transcriptional regulation

*In situ* and *in silico* expression analyses detected a peak of *ERF85* expression in phloem, cambial and expanding xylem (Figure 1). This agrees with the expression pattern observed for the closest homolog of *ERF85* in *A. thaliana*, *AtCRF4*, which is also highest expressed in phloem and xylem tissues [22]. Ectopic expression of *ERF85* throughout xylem cell expansion and SCW formation caused an increase in xylem fiber diameter, while simultaneously reducing SCW thickness and wood density in hybrid aspen (Figure 2). Interestingly, although the transcriptional induction of *ERF85* suggests that its function is regulated by ethylene, neither ethylene treatment itself [13, 14] nor ectopic expression of other ethylene-regulated *PtERFs* [15, 18] showed similar xylem cell phenotypes in hybrid aspen. Upon treatment with ACC, ERF85OE lines showed enhanced cambial activity and G-layer formation as previously observed in wild type trees [14], indicating that ERF85OE lines respond normally to ACC. A previous transcriptome study revealed that *ERF85* is induced by a 10h ACC application downstream of the ethylene signaling pathway [14]. We could not confirm this ACC-responsiveness using *pERF85::GUS* lines in our experiment. It should however be noted that the GUS-expression was rather weak under *in vitro* conditions (used for ACC treatments) as compared with greenhouse grown trees (Figure 1 and 2).

The differences in wood morphology and density in ERF85OE correlated with an induction of expansion zone associated genes and a repression of SCW-associated genes (Figure 4(e)) as well as with de-regulation of transcriptional networks in xylem cell expansion and SCW formation (Figure 5). Genes that affect cell elongation, like *CEL1* [33, 34] and *GAST1* [42], were upregulated by ectopic expression of *ERF85* (Figure 4(c)). In contrast, expression of genes involved in xylem maturation (e.g. *VNI2, XCP2*), lignin biosynthesis (e.g. *4CL2*, *CHS*) and SCW deposition (e.g. *IRX10*, *GH9C2*, *GH9B5*) were down-regulated in ERF85OE (Figure 4(c), Figure 5(d)). Two possible scenarios can explain this finding. Either ERF85 has a dual function and specifically induces genes of the cell expansion zone, while repressing those in the SCW formation zone, or the prolongated expression of expansion-associated genes leads to an indirect, delay in the onset of SCW gene expression. Furthermore, also a combination of both mechanisms may be at the origin of the observed phenotypes of enhanced fiber expansion and reduced SCW thickness in ERF85OE lines. It is widely accepted that TFs can function as repressors or activators depending on their target, entourage and interacting factors. Out of 1752 unique *A. thaliana* genes found among all homologs of ERF85-regulated DEGs, 995 have also been identified as direct targets of AtCRF4 in *A. thaliana* (Figure 6(a); Supplement Table S2) and those contain both up- and downregulated genes in our dataset. This favors the assumption that ERF85 could indeed have a dual regulatory function depending on its targets. On the other hand, induction of *CEL1* (Figure 4(c)) might trigger cell growth, while suppressing SCW thickening as seen in Arabidopsis [33]. It is likely that the phenotype is caused by a combined direct and indirect action of *ERF85*.

### 4.2 ERF85 might control xylem cell wall properties to allow cell expansion

In line with prolonged cell expansion and increased fiber diameter in ERF85OE, we also observed transcriptional changes in genes regulating cell wall properties, particularly pectin, required for cell expansion. Genes encoding enzymes with potential homogalacturonan backbone degradation function (*PtPG55* (M12), *PtPG2* (M5), *PtPG9* (M4)) were upregulated, indicative of increased pectin loosening that facilitates cell expansion, as for example shown for *PtPG28/43* homolog in Arabidopsis *AtPGX3* [36]). Modification of poplar *CEL1* expression has been shown to correlate with cell growth and cellulose content in suspension-cultured poplar [34]. These phenotypic responses were pronounced by the presence of auxin and sucrose [34]. Indeed, auxin is known to stimulate cell expansion by inducing cell wall loosening [43]. Hereby, auxin triggers demethylesterification of pectins, which can lead to enhanced wall extensibility in the presence of pectin degrading enzymes [44]. To gain cell wall rigidity in SCWs, auxin must be exported by auxin transporters like PIN3, which has been suggested to transport auxin in *Populus* stems [45]. In agreement with this fact, expression of *PtPIN3* is up-regulated during SCW formation (Figure 5(d)). Expression of *PtPIN3* and auxin signalling genes (e.g. *PtIAA4*, *PtIAA11*, *SAUR-like* genes) was down-regulated in transgenic ERF85OE trees, suggesting that ERF85 modifies auxin fluxes. The potential connection between ERF85 and polar auxin transport via PIN proteins agrees with previous reports that showed direct regulation of *AtPIN1* and *AtPIN7* by *AtCRF2* and *AtCRF6* in *A. thaliana* [31]. Thus, ERF85 could enhance cell expansion by transcriptional regulation of pectins, *CEL1* and auxin transport.

### 4.3 ERF85-mediated regulation of translation could affect the dynamics of cell growth in expanding xylem

Ectopic expression of *ERF85* induced the expression of all catalytical and regulatory centres of protein synthesis (Supplement Figure S3; Supplement Table S2). A recent study in *A. thaliana* revealed a new role of histone deacetylases (AtHD2B and AtHD2C) in transcriptional and posttranscriptional regulation of ribosome biogenesis [35]. Expression of two histone deacetylases *PtHD2A* and *PtHD2C* was strongly up-regulated in our ERF85OE lines (Figure 6(b), Supplement Table S2). Furthermore, gene regulatory network analysis also links expression of *ERF85*, *PtHD2C* and *PtGATAs* (*PtGATA15* and *PtGATA17L*; Figure 6(a)). In agreement with this, we identified the DRE-motif in the promoter of *PtGATA15* and *PtHD2C*, and a GCC-box in the promoter of *PtGATA17L*. Furthermore, *AtGATA17L* was identified as direct target of AtCRF4 in *A. thaliana* (Figure 6(a); Supplement Table S9; [38, 39]), suggesting that ERF-mediated transcriptional regulation of the gene network underlying xylem cell expansion might involve PtGATA15, PtGATA17L and PtHD2C as co-factors or potential second tier transcription factors. Future work will be required to identify direct targets of ERF85 and the molecular and functional importance of ERF85-regulated TFs.

## 5. Conclusions

Our study revealed ERF85 as regulator of transcriptional programs underlying xylem cell expansion and SCW formation. ERF85 stimulates fiber cell expansion and represses SCW formation probably through a combination of direct and indirect mechanisms. We therefore suggest that ERF85 acts as a control point for the regulation of transcriptional programs during these two developmental phases of secondary xylem development.

---

Supplement Figure S1: The *LMX5* promoter extends expression of *ERF85* in *ERF85OE* beyond xylem expansion.

Supplement Figure S2: Ectopic expression of *ERF85* does not affect vessel diameter.

Supplement Figure S3: Summary of enriched GO-terms among down- and up-regulated DEGs of ERF85OE.

Supplement Table S1: Primer used in this study

Supplement Table S2: DEGs identified in ERF85OE

Supplement Table S3: Enriched GO terms in down-regulated DEGs

Supplement Table S4: Enriched GO terms in up-regulated DEGs

Supplement Table S5: Gene modules identified according to AspWood

Supplement Table S6: GO terms enriched in each gene module

Supplement Table S7: Representative genes shown for each gene module

Supplement Table S8: Statistics for promoter analysis

Supplement Table S9: TF network regulated by *ERF85*

## Author Contributions

CS analyzed and quantified the SCW pictures, analyzed the RNA-Seq data and performed bioinformatic and statistical analyses. BW cloned the pERF85::GUS construct, generated and selected transgenic trees and performed the GUS staining. JV and JK carried out the wood density measurements. ND helped with the RNA-Seq analysis. TRH performed the motif analysis. JF constructed the poplar experiments and acquired microscopy pictures for the wood anatomy analysis of ERF85OE. JF designed and initiated the project and experiments. CS, BW, JF and HT analyzed and discussed the data. The manuscript was written by CS and JF with contributions from all co-authors.

## Data Availability Statement

Raw files can be downloaded from the European Nucleotide Archive (ENA) under PRJEB35743. Data quality summary and scripts used in this study were deposited to https://github.com/carSeyff/ERF85OE.

## Acknowledgments

We thank Peter Grones for help with picture analysis, Tuomas Puukko for technical help in laboratory analyses, Pekka Lönnqvist and Leena Grönholm in greenhouse experiments, the Umeå Core Facility Electron Microscopy for generating ultrathin sections. Computations were performed on resources provided by the Swedish National Infrastructure for Computing (SNIC) at UPPMAX.

