## Supplement_Figures_S1-S3 for "*Populus* ERF85 balances xylem cell expansion and secondary cell wall formation in hybrid aspen"

### Slide 1
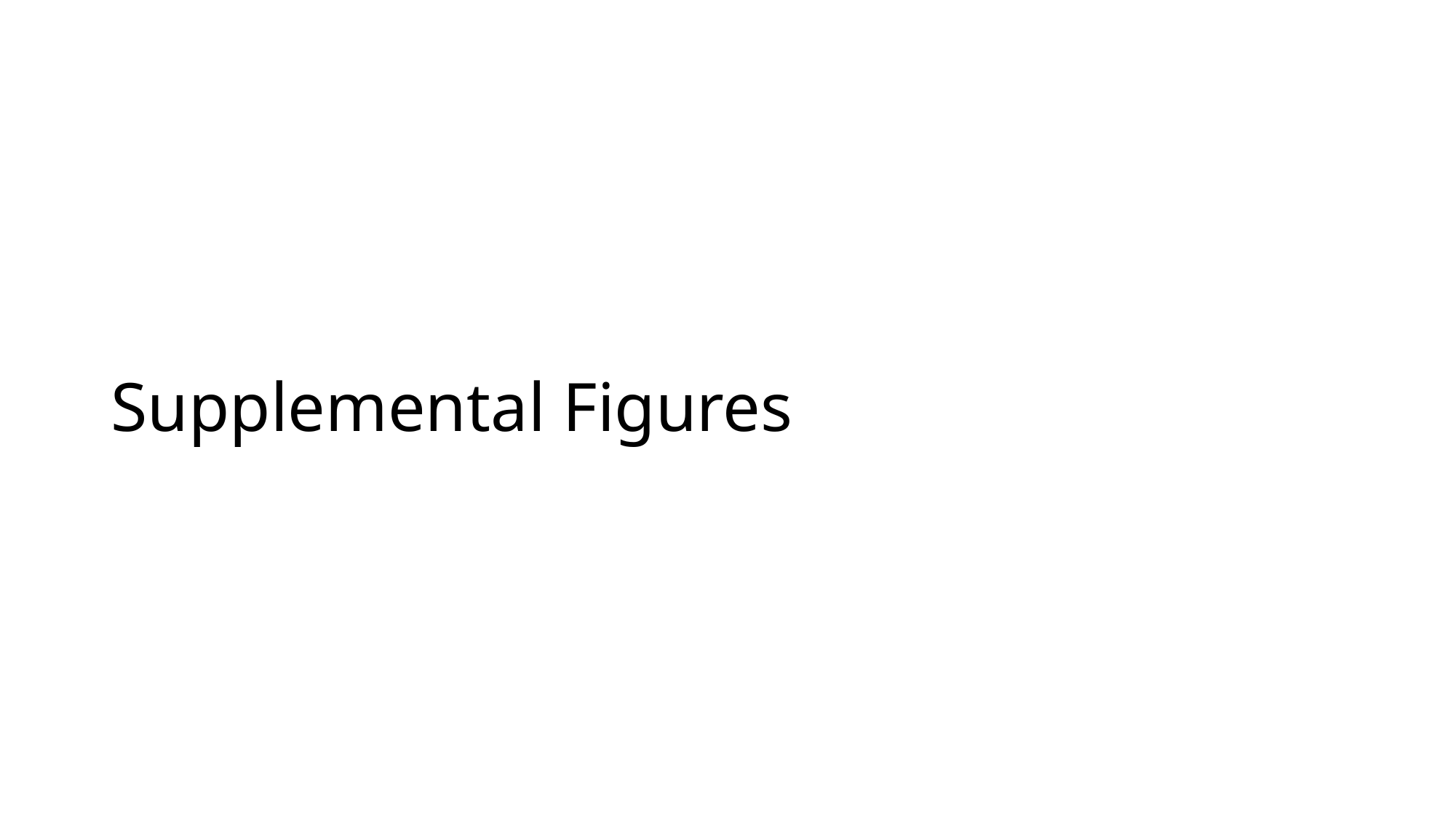

#### Supplemental Figures

### Slide 2
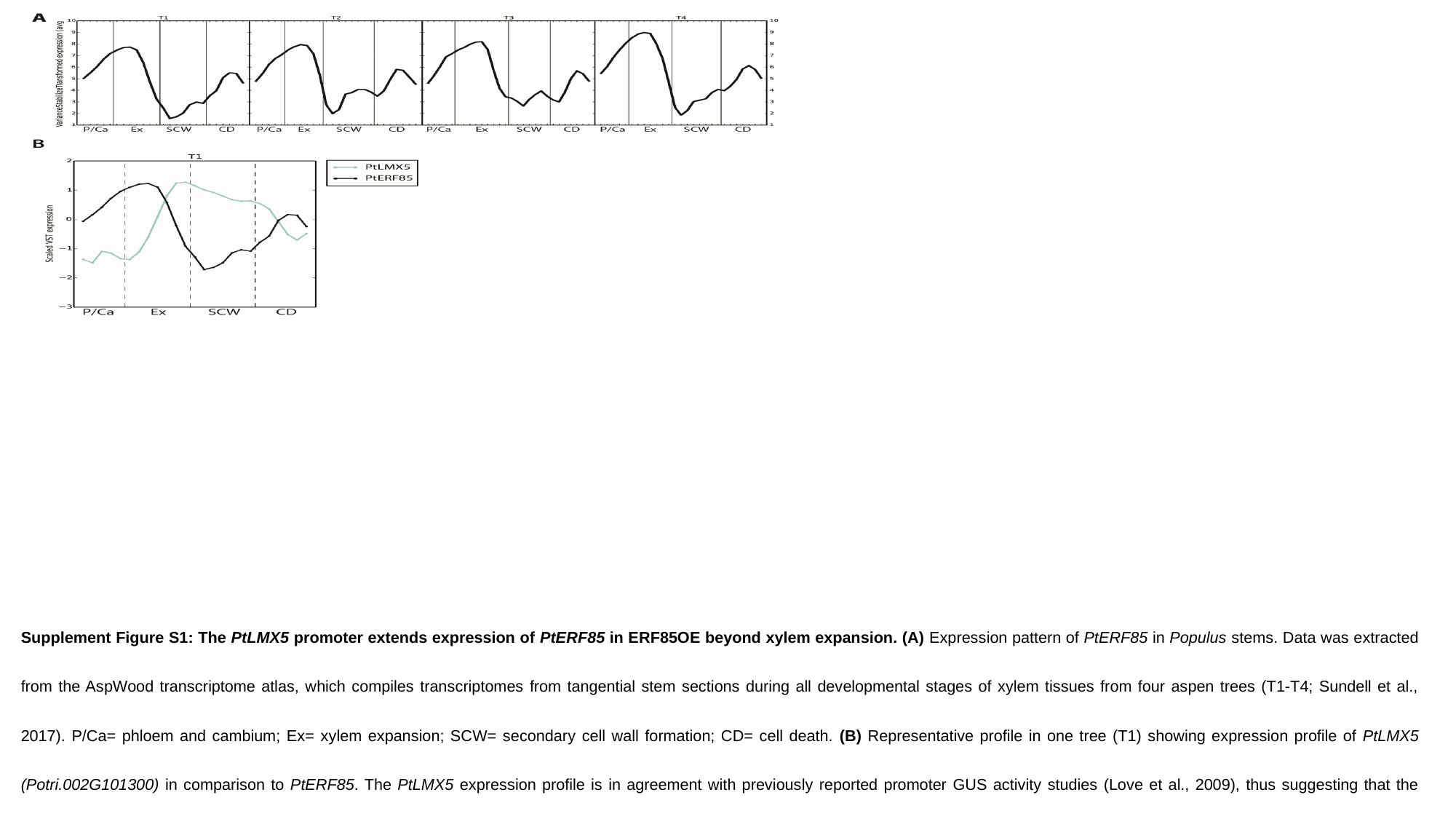

^Supplement Figure S1: The^ ^PtLMX5^ ^promoter extends expression of^ ^PtERF85^ ^in ERF85OE beyond xylem expansion. (A)^ ^Expression pattern of^ ^PtERF85^ ^in^ ^Populus^ ^stems. Data was extracted from the AspWood transcriptome atlas, which compiles transcriptomes from tangential stem sections during all developmental stages of xylem tissues from four aspen trees (T1-T4; Sundell et al., 2017). P/Ca= phloem and cambium; Ex= xylem expansion; SCW= secondary cell wall formation; CD= cell death.^ ^(B)^ ^Representative profile in one tree (T1) showing expression profile of^ ^PtLMX5 (Potri.002G101300)^ ^in comparison to^ ^PtERF85. The^ ^PtLMX5^ ^expression profile is in agreement with previously reported promoter GUS activity studies (Love et al., 2009), thus suggesting that the^ ^PtLMX5^ ^promoter extends^ ^PtERF85^ ^expression into the SCW forming zone. Lines represent scaled and smoothed VST expression values obtained from the AspWood database.^

### Slide 3
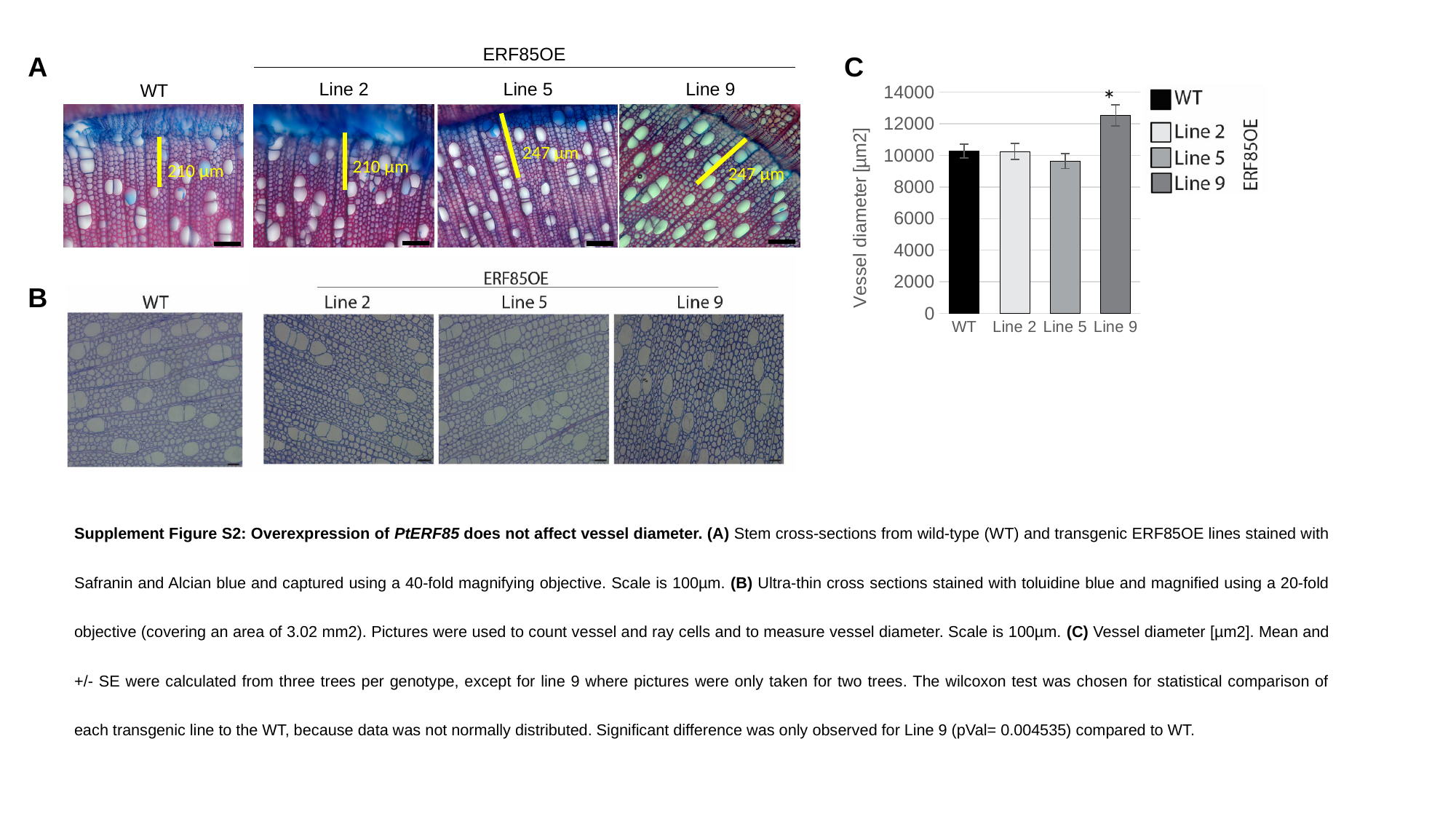

ERF85OE
A
C
Line 2
Line 5
Line 9
WT
###### Chart
| Category | Mean |
|---|---|
| WT | 10270.26 |
| Line 2 | 10251.41 |
| Line 5 | 9643.778 |
| Line 9 | 12533.6 |
| Line 12 | 10697.3 |
| Line 13 | 11286.35 |*
247 µm
210 µm
210 µm
247 µm
B
^Supplement^ ^Figure^ ^S2: Overexpression of^ ^PtERF85^ ^does not affect vessel diameter. (A)^ ^Stem cross-sections from wild-type (WT) and transgenic ERF85OE lines stained with Safranin and Alcian blue and captured using a 40-fold magnifying objective. Scale is 100µm.^ ^(B)^ ^Ultra-thin cross sections stained with toluidine blue and magnified using a 20-fold objective (covering an area of 3.02 mm2). Pictures were used to count vessel and ray cells and to measure vessel diameter. Scale is 100µm.^ ^(C)^ ^Vessel diameter [µm2]. Mean and +/- SE were calculated from three trees per genotype, except for line 9 where pictures were only taken for two trees. The wilcoxon test was chosen for statistical comparison of each transgenic line to the WT, because data was not normally distributed. Significant difference was only observed for Line 9 (pVal= 0.004535) compared to WT.^

### Slide 4
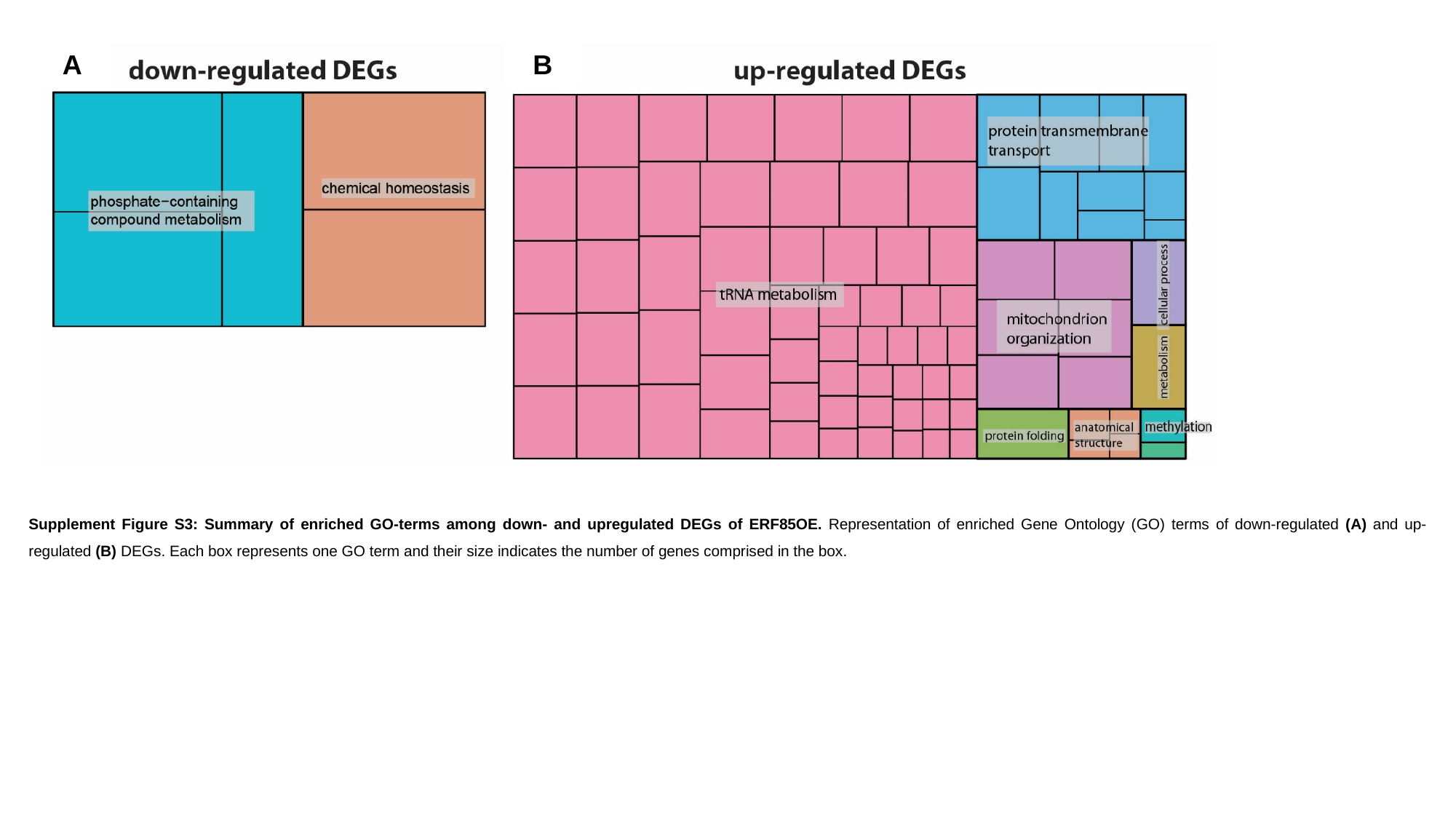

A
B
Supplement Figure S3: Summary of enriched GO-terms among down- and upregulated DEGs of ERF85OE. Representation of enriched Gene Ontology (GO) terms of down-regulated (A) and up-regulated (B) DEGs. Each box represents one GO term and their size indicates the number of genes comprised in the box.
